# The therapeutic potential of D-Serine in reducing expression of the cytopathic genotoxin colibactin

**DOI:** 10.1101/2022.03.08.483572

**Authors:** Jennifer C Hallam, Nicky O’Boyle, Natasha C A Turner, Min Tang-Fichaux, Eric Oswald, Andrew J Roe

## Abstract

Som*e Escherichia coli* strains belonging mainly to the B2 phylogroup harbour the *pks* island, a 54 kb genomic island encoding the biosynthesis genes for a genotoxic compound named colibactin. In eukaryotic cells, colibactin can induce DNA damage, cell cycle arrest and chromosomal instability. Moreover, production of colibactin has been implicated in the development of colorectal cancer. In this study, we demonstrate the inhibitory effect of D-Serine on the expression of the *pks* island in two colibactin-producing strains, CFT073 and Nissle 1917, and determine the implications for cytopathic effects on host cells. To investigate the specificity of the inhibitory effect of D-Serine, we also tested a comprehensive panel of proteinogenic L-amino acids and corresponding D-enantiomers for their ability to modulate *clbB* transcription using RT-qPCR. Several D-amino acids exhibited the ability to inhibit expression of *clbB*, with D-Serine exerting the strongest repressing activity (3.81-fold in CFT073; 3.80-fold in Nissle 1917) and thus, we focussed additional experiments on D-Serine. To investigate the cellular effect, we investigated if repression of colibactin by D-Serine could reduce the cytopathic responses normally observed during infection of HeLa cells with *pks*^+^ strains. Levels of γ-H2AX (a marker of DNA double strand breaks) were reduced 2.75-fold in cells infected with D-Serine treatment. Moreover, exposure of *pks*^+^ *E. coli* to D-Serine during infection caused a reduction in cellular senescence that was observable at 72 h post infection. The recent finding of an association between *pks*-carrying commensal *E. coli* and CRC, highlights the necessity for the development of colibactin targeting therapeutics. Here we show that D-Serine can reduce expression of colibactin, and inhibit downstream cellular cytopathy, illuminating its therapeutic potential to prevent colibactin-associated disease.

## Introduction

The gastrointestinal tract is a complex environment, where the composition and abundance of metabolites and signalling molecules can be affected by the host diet and physiology, the microbiota, and invading pathogens [1]. How bacteria sense and respond to these signals, is crucial for their survival and successful colonization within the host. Most of the colonizers are commensal organisms which live mutualistically, however, certain pathogenic strains can outcompete resident bacteria by utilizing host metabolites. Indeed, competition in this environment goes beyond contending for food as metabolites have also been shown to influence gene expression and drive evolutionary change amongst bacteria residing in the gut [2].

The composition of the human diet greatly influences the availability of metabolites within the gastrointestinal environment, as many indigestible components provide substrates for the resident microflora. An often-overlooked group of metabolites are the D-amino acids, of which the average dietary consumption is estimated at 100 mg/d [3]. D-amino acids are found naturally in foods such as fruits and vegetables [4], however, higher concentrations are more commonly associated with fermented and processed foods, including aged cheeses and breakfast cereals [3]. D-amino acids are also an important source of energy for the resident microflora. Indeed, bacteria can utilize these for essential processes including supporting growth, regulating spore germination, and in cell wall synthesis, where D-Alanine and D-Glutamate are routinely found as components of the peptidoglycan sacculus [5]. Furthermore, D-amino acids have been found to have a profound effect on gene expression and studies have shown they can manipulate the expression of virulence genes in diverse bacteria [6–8]. The concentrations of these D-amino acids are often dependent on the site within the host. For example, D-Serine is found at 1 μM in the gut and at 1 mM in the urine [9], a substantial difference that we believe could play a role in regulating niche-specificity [10]. In order to metabolize these molecules, bacteria must possess specialized catabolic enzymes [11–16]. In the case of D-Serine, certain bacteria encode a specialized tolerance locus, enabling them to exploit D-Serine as a carbon source, facilitating colonization at nutrient deficient extraintestinal sites. The uropathogenic *Escherichia coli* (UPEC), commonly possess this capability. UPEC strains can traverse through the gastrointestinal tract and colonize the bladder, where carbon sources are scarce, but D-Serine concentrations are high [17,18]. UPEC, and many other *E. coli* that belong to the B2 phylogroup, commonly encode a D-Serine tolerance locus, *dsdCXA*. This locus encodes a D-Serine deaminase (DsdA), a D-Serine inner membrane transporter (DsdX) and an essential LysR-type transcriptional regulator (LTTR) (DsdC) that regulates the system [16]. Indeed, the role of *dsdCXA* is to prevent UPEC strains from succumbing to inhibitory concentrations of D-Serine in the urinary tract, by converting the substrate to ammonia and pyruvate [9,16,19]. While there have been conflicting reports surrounding the role of D-Serine in regulating virulence in UPEC [9,18,20], recent transcriptome analysis has demonstrated that D-Serine exhibits the ability to modulate the expression of genes beyond the *dsdCXA* locus in both *dsd*- and *dsd*+ strains [21].

Recently, we described the transcriptional response to D-Serine in strains from three distinct *E. coli* pathotypes; enterohaemorrhagic *E. coli* (EHEC), neonatal meningitis associated *E. coli* (NMEC) and UPEC [21]. Strikingly, the results revealed a unique transcription profile for each pathotype with not a single differentially expressed gene shared between the strains. Interestingly, the transcriptional response in UPEC highlighted differential expression in a cluster of genes involved in a non-ribosomal peptide synthetase pathway. This peptide synthetase pathway, termed the *pks* island, was first described in 2006, where *E. coli* encoding this 54 kb genomic island were implicated with inflicting a genotoxic insult on eukaryotic cells [22]. The *pks* island is comprised of 19 genes which encode the machinery for biosynthesis and transportation of colibactin, the peptide-polyketide hybrid compound responsible for exerting genotoxic activity [23]. The *pks* island was first identified in extraintestinal *E. coli* (ExPEC) strains, with colibactin described as a *bona fide* virulence factor in several studies [24–27]. Carriage of the *pks* island has also been described in around 34% of commensal *E. coli* strains belonging to the B2 phylogenetic group [23,27,28], and in other members of the *Enterobacteriaceae* including *Citrobacter koseri*, *Klebsiella pneumoniea* and *Enterobacter aerogenes* [23].

Recently, there has been a dramatic increase in interest in colibactin research, largely due to the identification of an association between colibactin activity and the development of colorectal cancer (CRC). This association arises from the ability of colibactin to cause DNA double-strand breaks (DSBs), DNA crosslinks and chromosome instability in eukaryotic cells [29–32]. *E. coli* harbouring the *pks* island have been isolated from biopsy specimens of CRC patients and have been found to be capable of establishing persistent colonization, inducing inflammation, and triggering tumour growth in these tissues [33–35]. Indeed, recent whole genome analysis of *pks*^+^ infected organoids revealed a distinct mutational signature in adenine rich residues. This mutational signature was also observed in CRC tissues, and it was confirmed that the mutation was specific to colibactin exposure [36,37]. CRC is the third most frequently diagnosed cancer [38] and attributes to around 610,000 deaths per year worldwide [35]. Therefore, colibactin-producing *E. coli* represent an urgent public health matter. Here, we compared the effects of a comprehensive panel of naturally-occurring amino acids – comprising the 20 proteinogenic L-amino acids and their corresponding D-enantiomers – on the expression of colibactin synthesis genes. Consistent with our recent study [21], we found that exposure to D-Serine induced a significant downregulation of the *pks* gene cluster. Further, we tested the effects of D-Serine during infection of HeLa cells and found that treatment of *pks* harbouring *E. coli* resulted in a dampening of the genotoxic effect exerted upon eukaryotic cells. Considering this research, we propose D-Serine as a novel therapeutic to control expression of colibactin in *pks*-encoding *E. coli* strains.

## Materials and Methods

### Bacterial strains, plasmids and cultures

The bacterial strains, plasmids and oligonucleotides used in this study are listed in Table S1 and Table S2. Bacteria were routinely grown at 37°C in Luria broth (LB [Miller’s recipe]) before diluting 1/100 into the appropriate medium for experiments or growth analysis. Chloramphenicol was used when appropriate at a concentration of 25 µg/ml. All preparations of M9 minimal medium (Sigma Aldrich; cat# M6030) were supplemented with 0.4% (*w/v*) glucose unless otherwise stated. For HeLa cell infection experiments, bacteria grown overnight were inoculated in prewarmed MEM-HEPES (Sigma Aldrich; cat# M7278) −/+ 1 mM D-Serine and incubated at 37°C, 200 RPM for 4.5 h. All growth media, antibiotics and chemicals were purchased from Sigma Aldrich unless stated otherwise.

### Assessment of DNA crosslinking activity

The assay was performed as previously described [39]. Briefly, linearized plasmid DNA was generated by digestion of pUC19 plasmid with BamHI (New England Biolabs). For bacteria-DNA interactions, bacteria were inoculated 1:20 from overnight cultures into 10 mL M9 Minimal Media alone or supplemented with 1 mM D-Serine and grown for 1.5 h. After reaching an OD_600 nm_ of ~0.6, 1 × 10^6^ CFU was inoculated in 100 μL M9 Minimal media alone or supplemented with 1 mM D-Serine and incubated statically at 37°C for 5 h. Following, cultivation, cells were harvested by centrifugation and the media was removed. Cells were resuspended in sterilised nuclease-free water. Then, a mixture of 450 ng of linearized DNA and 1 mM EDTA was added and samples were incubated for a further 40 min. Bacteria were pelleted by centrifuging samples at 5,000 × g for 5 min, then the DNA present in the supernatants was purified using the PCR Purification kit (Qiagen) according to the manufacturer’s instructions. A denaturing 1% agarose gel was prepared in a 100 mM NaCl and 2 mM EDTA solution (pH 8.0), then the gel was soaked overnight in an alkaline running buffer solution (40 mM NaOH and 1 mM EDTA, pH ~12). For each sample, 100 ng of DNA was loaded on to the denaturing agarose gel and then the gel was run at 1 V/cm for 45 min and then for 2 h at 2V/cm. The gel was neutralized in a 100 mM Tris pH 7.4 buffer solution containing 150 mM NaCl, that was frequently changed, for a total of 45 min. The gel was stained with GelRed and DNA was revealed with UV exposure using the ChemiDoc Imaging System (BioRad).

### HeLa cell culture, infection and examination of cellular senescence

For bacterial infections, overnight LB cultures of bacteria were cultured in prewarmed MEM-HEPES −/+ 1 mM D-Serine and cultured for 4.5 h at 37°C with 200 RPM agitation. Bacterial suspensions were diluted to OD _600 nm_ of 0.1, before serially diluting and spot plating on LB plates to confirm appropriate cell density. HeLa cells were routinely cultured in DMEM (ThermoFisher Scientific; cat# 61965026) with 10% (*v/v*) foetal calf serum (FCS) at 37°C, in a 5% CO_2_ incubator and were maintained by serial passage. For infection experiments, 4 × 10^4^ cells/well were seeded on 13 mm glass coverslips pre-coated with collagen (Millipore; cat# 08-115) as per the manufacturer’s instructions. After 24 h and immediately prior to infection, the HeLa cells were washed with DPBS (ThermoFisher Scientific, cat# 14190086) and medium was replaced with MEM-HEPES −/+ 1 mM D-Serine. Bacteria were added to each coverslip at a multiplicity of infection of 400 (200 μl of 0.1 OD_600 nm_ suspension) and infected for 4 h. The cells were washed twice with DPBS 4 h after inoculation, then cells were replenished with DMEM containing 10% (v/v) FCS and 50 μg/ml gentamicin (Sigma Aldrich, cat# G1397) and incubated for 72 h at 37°C, 5% CO_2_. Next, the cells were fixed in 4% (*w/v*) paraformaldehyde at room temperature for 15 min. The cells were then permeabilized with 0.1% (*v/v*) Triton X-100 in DPBS for 5 min. After two washes, each coverslip was stained with 0.2 U Phalloidin-Alexa Fluor 555 (Invitrogen, cat# A34055) for 1 h at room temperature. The cells were washed twice with DPBS before the coverslips were mounted on to a glass slide with 4 μl Vectashield with DAPI (Vector Laboratories, cat# H-1200) and sealed with clear nail polish. Images were acquired using a Zeiss AxioImager M1 and images were processed by deconvolution using Zen 2.3 Pro software (Zeiss). The area of each cell was measured using a pipeline developed on CellProfiler [40]. Briefly, sample images were acquired at 10X magnification to allow for >100 cells to be captured per image. Image files were uploaded to the CellProfiler workspace and analysis was performed for images taken from 3 replicate experiments.

### Immunofluorescence analysis of H2AX phosphorylation

HeLa cells were seeded on 13 mm coverslips and infected as described above. Following 4 hours infection cells were washed twice with DPBS and fixed in 4% (*w/v*) paraformaldehyde for 15 min at room temperature. The cells were permeabilized with 0.1% Triton X-100 and then blocked with 1X Phosphate-Buffered Saline, 0.1% Tween^®^ 20 (PBST) + 10% normal goat serum (Sigma Aldrich, cat# NS02L) for 1 h at room temperature. Next cells were incubated with rabbit monoclonal anti γ-H2AX antibodies (Cell Signalling, cat# 5438S) diluted 1:100 in blocking solution and incubated for 1 h at room temperature. The tissues were washed three times with DPBS, then a fluorescent secondary antibody, Alexa Fluor 555 Goat anti-rabbit IgG (Invitrogen, cat# A32732) diluted 1:400 in blocking solution was applied and incubated for 1 h in the dark at room temperature. Following incubation, tissues were washed three times with DPBS, then coverslips were mounted to a glass slide with 4 μl Vectashield with DAPI and sealed with clear nail polish. Nuclear foci were visualized using a Zeiss LSM 880 confocal microscope (Zeiss).

### Flow cytometry analysis of H2AX phosphorylation

Twenty-four well tissue culture plates were collagen-coated, seeded with 10^6^ cells/well and infected as described above. The cells were collected by trypsinization 4 h post-infection and washed in DPBS. The cells were then collected by centrifugation and resuspended in a live/dead stain (eFluor 780 [eBioscience, cat# 65-0865-14]) and incubated for 20 min on ice. The cells were washed in excess Stain Buffer (BD Biosciences, cat# 55456) and FC receptor block (DPBS + 10% FCS) was applied before incubating for a further 20 min on ice. Next, the cells were fixed with Cytofix (BD Biosciences, cat# 554655) for 15 min at 37°C and permeabilized with Perm Buffer (BD Biosciences, cat# 558050) for 30 min on ice, before purified mouse anti γ-H2AX (BD Biosciences, cat# 560443) diluted 1:200 in BD Stain Buffer was applied and cells were incubated for 1 h at room temperature. A multichromatic-conjugated secondary antibody, goat anti-mouse IgG (BD Biosciences, cat# 550589) diluted 1:1000 in Stain Buffer was applied and cells were incubated for 1 h at room temperature, keeping samples protected from the light. Cells were eventually suspended in BD Stain Buffer and filtered with a 70 μm filter. Cells were analysed using the BD FACSAria (BD) and the data was analysed using FloJo software. Analysis of the stained cell populations was performed by gating on single, live cells.

### Western Blot analysis of H2AX phosphorylation

HeLa cells were infected with Nissle 1917 (MOI = 400) for 4 h then treated with gentamicin for a further 4 h. The tissues were washed and lysed directly in the cell culture well by applying 100 μl of 1X SDS sample buffer and incubating for 5 min then wells were scraped to release attached cells, cell lysates were immediately stored on ice. The cell lysates were heated for 10 min at 90°C, and aliquots were stored at −20°C. Proteins were separated on 4-12% Bis Tris Gel by SDS-PAGE (Invitrogen, cat# NP0321) and transferred to a nitrocellulose membrane (FisherScientific, cat# 88018). Blocking was performed using 5% skimmed milk powder for 1 h in PBST. The membrane was then incubated with anti-γ H2AX primary antibody (Cell Signalling Technologies, cat# 5438S) diluted 1:100 in 5% BSA-PBST and incubated overnight at 4°C. The membrane was then washed three times with PBST for 10 min before being incubated for 1 h with anti-rabbit horseradish peroxidase (HRP)-conjugated secondary antibody (Invitrogen, cat# 656120) diluted 1:2500 in PBST. The membrane was again washed three times with PBST for 10 min. Bound secondary HRP-labelled antibodies were revealed with SuperSignal™ West Femto maximum sensitivity substrate (ThermoFisher Scientific, cat# 34096) and analysed with the C-DiGit^®^ blot scanner (LI-COR). Membranes were stripped with mild stripping buffer and incubated with the primary antibody H2AX (Cell Signalling Technologies, cat# 25955) and detected with the secondary antibody as described above. To control for sample loading, membranes were probed for β-Tubulin (Abcam, cat# ab6046). Proteins were quantified with Image Studio Lite (Licor) and normalized in relation to the β-Tubulin level.

### RNA Extraction

For screening of amino acids capable of modulating expression of *clbB*, bacteria were cultured in M9 Minimal Media supplemented with amino acids at 1 mM final concentration for 5 h. The cells were harvested by centrifugation before resuspension in two volumes of RNAprotect Bacteria Reagent (Qiagen, cat# 76506). After a 5 min incubation at room temperature, cells were harvested, and RNA extractions were carried out using PureLink RNA Mini Kit (Thermofisher Scientific, cat# 12183018A) according to manufacturer’s instructions. Contaminating DNA was removed by TurboDNase (ThermoFisher Scientific, cat# AM2238) treatment, followed by extraction in phenol-chloroform-isoamyl alcohol (Sigma Aldrich, cat# P2069) and ethanol precipitation at −80°C overnight. RNA was collected by centrifugation before washing with 70% ethanol and resuspending in nuclease-free water.

### Synthesis of cDNA and RT-qPCR

Ten nanograms of DNA-free total RNA, extracted as described above, was used as a template to prepare 10 μl cDNA using LunaScript^®^RT SuperMix Kit (New England Biolabs, cat# E3010L) according to manufacturer’s instructions. Luna^®^ Universal qPCR Master mix (New England Biolabs, cat# M3003L) was employed for RT-qPCR according to manufacturer’s recommendations (Initial denaturation 95°C, 1 min; denaturation 95°C, 15 sec; extension 60°C, 30 sec; 39 cycles). Technical duplicate 20 μl reactions were carried out with 1 μl volumes of cDNA as template and triplicate biological samples of cDNA being analysed. Expression relative to the untreated control was calculated as 2^−ΔΔct^ with [41] *gapA* amplification being employed as a housekeeping control.

### Assessment of *clbB* expression using promoter-*gfp* fusion reporter

The promoter region upstream of *clbB* was amplified using the P*clbB*-Fw and P*clbB*-Rev primers Table S2, before digesting with BamHI/KpnI and cloning in frame upstream of green fluorescent protein (*gfp*) in pAJR70 [42]. Bacteria were cultured in MEM-HEPES for 4 h before recording OD_600 nm,_ and fluorescence using a BMG Fluostar plate reader. Relative fluorescence was determined by subtracting the fluorescence intensity of an empty vector control from each sample and dividing the corrected fluorescence intensity by OD_600 nm_. Data presented are from three replicate experiments.

### Construction of isogenic ∆*dsdC* Nissle 1917 mutant strain

A Nissle 1917 mutant lacking *dsdC* was constructed as described elsewhere [43]. Briefly, Nissle 1917 WT was transformed with pKD46. A single colony was cultured at 37°C in LB supplemented with 100 µg/ml ampicillin and 100 mM L-Arabinose to an OD_600 nm_ of 0.4. The cells were then washed and resuspended three times with ice-cold distilled water. A linear deletion fragment was prepared by amplifying the chloramphenicol resistance cassette from pKD3 with oligonucleotides bearing 50 bp 5’-end flanking regions homolgous to the 50 bp regions immediately upstream and downstream of *dsdC*, Table S2. One microgram of PCR product (phenol-chloroform extracted, before ethanol precipitation and resuspension in 10 μl nuclease-free water) was electroporated at 2500 V into an aliquot of competent Nissle 1917 cells using an Eppendorf Eporator. Insertional mutants grown under chloramphenicol selection were verified by PCR using check primers, Table S2. Resistance cassettes were removed by expression of FLP-recombinase under transient temperature shift to 42°C after transformation of insertional mutants with pCP20. Excision of the resistance cassette was confirmed by PCR.

### Statistical analysis

Statistical significance was assessed by unpaired two-tailed Student’s *t*-test or ordinary one-way ANOVA, as indicated. Tests were carried out using GraphPad Prism 8.

## Results

### Colibactin biosynthesis is downregulated in response to D-serine

The host metabolite D-Serine has been shown to selectively affect the expression of virulence factors in *E. coli* pathotypes that do not possess the intact *dsdCXA* locus [10]. However, little is known of the effects on gene expression in pathotypes found in D-Serine rich environments such as the bladder. Therefore, this prompted an investigation into how exposure to D-Serine implicated gene expression in *E. coli* that encoded the complete *dsdCXA* locus. Previous work examining the transcriptome of CFT073 exposed to D-Serine revealed many significant differentially expressed genes throughout the genome, but of particular interest was the downregulation of several genes encoding proteins involved in the synthesis of the genotoxin colibactin [21]. Comparison of read data from D-Serine treated (red) and untreated control (black) CFT073, indicate that treatment results in a downshift in expression of several colibactin synthesis genes (Fig 1A). The log_2_-transformed fold changes observed are indicated in Fig 1B. The most significant reductions were identified for genes *clbB, I, G, C, H* and *K* respectively. Full details for fold changes and *P*-values can be found in Table S3. Interestingly, these genes belong to four (*clbB*, *clbC-G*, *clbH* and *clbI-N*) of seven putative transcriptional units of the *clb* locus [44] and encode enzymes essential for production of precolibactin, the precursor to cytotoxic colibactin [45], suggesting D-Serine can interfere with the manufacture of active colibactin in CFT073.

**Fig 1.**
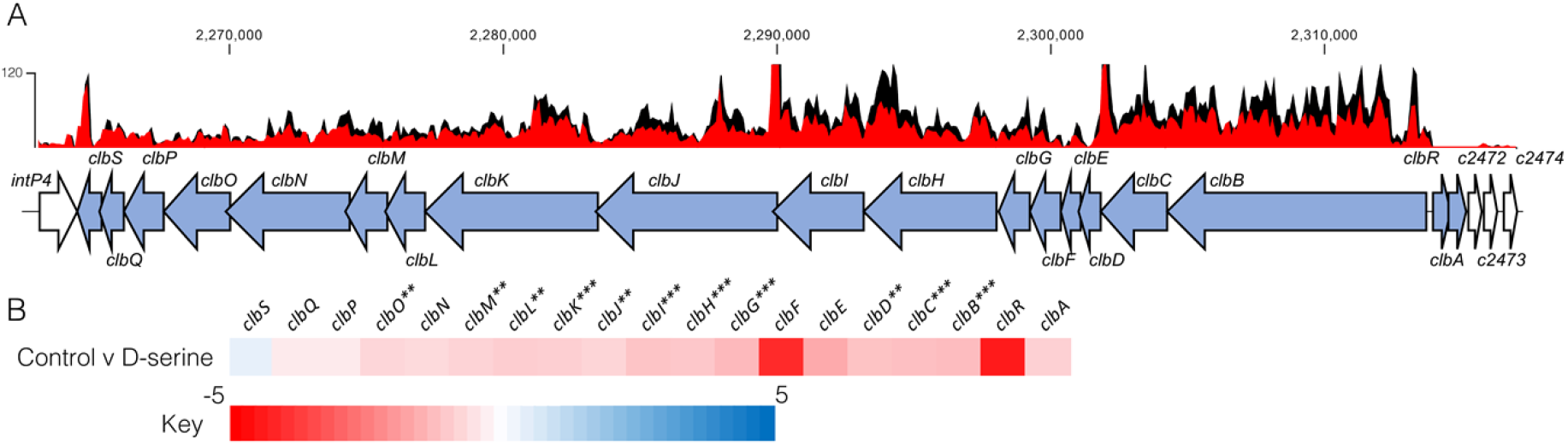
Transcriptome analysis of CFT073 reveals downregulation of the colibactin biosynthesis operon in response to D-Serine. Using read data from our previous transcriptomic study [21], we constructed track maps of the *clb* locus in CFT073 in the presence and absence of D-Serine. (**A**) Read density in the colibactin biosynthesis locus for representative samples of the untreated control (black) and D-Serine treated (red) CFT073 is indicated. Read tracks were normalized, exported from CLC Genomics Workbench and overlayed. Genomic coordinates are displayed above the read tracks and the corresponding genes within the colibactin biosynthesis operon (blue) beneath. (**B**) Heat map indicating the EdgeR calculated log_2_ relative fold changes for each gene in the colibactin biosynthesis locus with corresponding colour key beneath. False discovery rate-corrected *P* values are indicated with significantly differentially expressed genes with *, ** and *** indicating *P* < 0.05, 0.01 and 0.001, respectively.

### Expression of the genotoxin colibactin is affected by D-and L-amino acids in both CFT073 and Nissle 1917

We next investigated if the downregulation of the colibactin genes was a unique property of D-Serine, by comparing a comprehensive panel of L- and D-amino acids under the same growth conditions. Real-time quantitative polymerase chain reaction (RT-qPCR) was used to determine expression of the *pks* encoded gene *clbB*; chosen as it encodes a hybrid non-ribosomal peptide synthetase/type I polyketide synthase (NRP-PKS) that is integral to colibactin synthesis [46]. Exposing CFT073 to L-Aspartic acid, L-Isoleucine, and L-Selenocysteine led to the most significant decrease in *clbB* expression; reductions of 7.52, 8.84, and 6.17-fold were observed, respectively (Fig 2A). In addition, expression of *clbB* was significantly reduced in 9 of 20 D-amino acids tested, with the most significant observations recorded for expression of *clbB* in D-Cysteine (0.11) and D-Serine (0.32), *P* = 0.009 and 0.006, respectively (Fig 2B). Interestingly, of the seven L-amino acids showing a significant reduction, only the D-enantiomer of Isoleucine significantly changed expression of *clbB* in CFT073, highlighting the distinction in responses to L- and D-amino acids. Next, we selected some of the amino acids which displayed a significant effect on *clbB* expression in CFT073, and tested them with the commensal *E. coli* strain, Nissle 1917. This strain was chosen based on its commensal origin and clinical significance as a probiotic [47,48]. Furthermore, it has successfully been used in the characterization of the colibactin-associated phenotype observed in HeLa cells and recently the genotoxicity of Nissle 1917 was demonstrated in the gut lumen of mice [49,50], while CFT073 is precluded from such analysis as the cytopathic effects of colibactin are often confounded by haemolysin activity [51]. In response to the L- and D-amino acids tested (Fig 2C), significant downregulation of *clbB* in Nissle 1917 was observed for 7 of 17 amino acids tested. Interestingly, L-Selenocysteine yielded the largest decrease, with *clbB* expression reduced 5.64-fold compared to the control (*P* = 0.029). However, it should be noted that growth of both CFT073 and Nissle 1917 was strongly inhibited in the presence of L-Selenocysteine, likely accompanied by stress-associated transcriptomic perturbations that may have indirectly altered *clbB* expression in a not observed with other amino acids. The second most significant decrease was observed for D-Serine where a 3.81-fold reduction was observed, (*P* = 0.036). Relative expression and *P*-values for all amino acids tested can be found in Table S4 for CFT073, and S5 for Nissle 1917. Our RT-qPCR results corroborated the findings of our previous RNA-Seq study [21] and confirmed that D-Serine repressed expression of the colibactin synthesis genes in CFT073. In addition, D-Serine was shown to suppress *clbB* expression in Nissle 1917. Before testing the effects of D-Serine during infection of host cells, expression of *clbB* was measured by *gfp* reporter assay on cultures grown in MEM-HEPES tissue culture media. A similar response to D-Serine was observed in both strains, with *clbB* expression reduced by 2.00 and 1.75-fold in CFT073 and Nissle 1917, respectively (Fig 2D). These data suggest that both L- and D-amino acids can affect the expression of colibactin but most notably, the host metabolite D-Serine has a profound effect.

**Fig 2.**
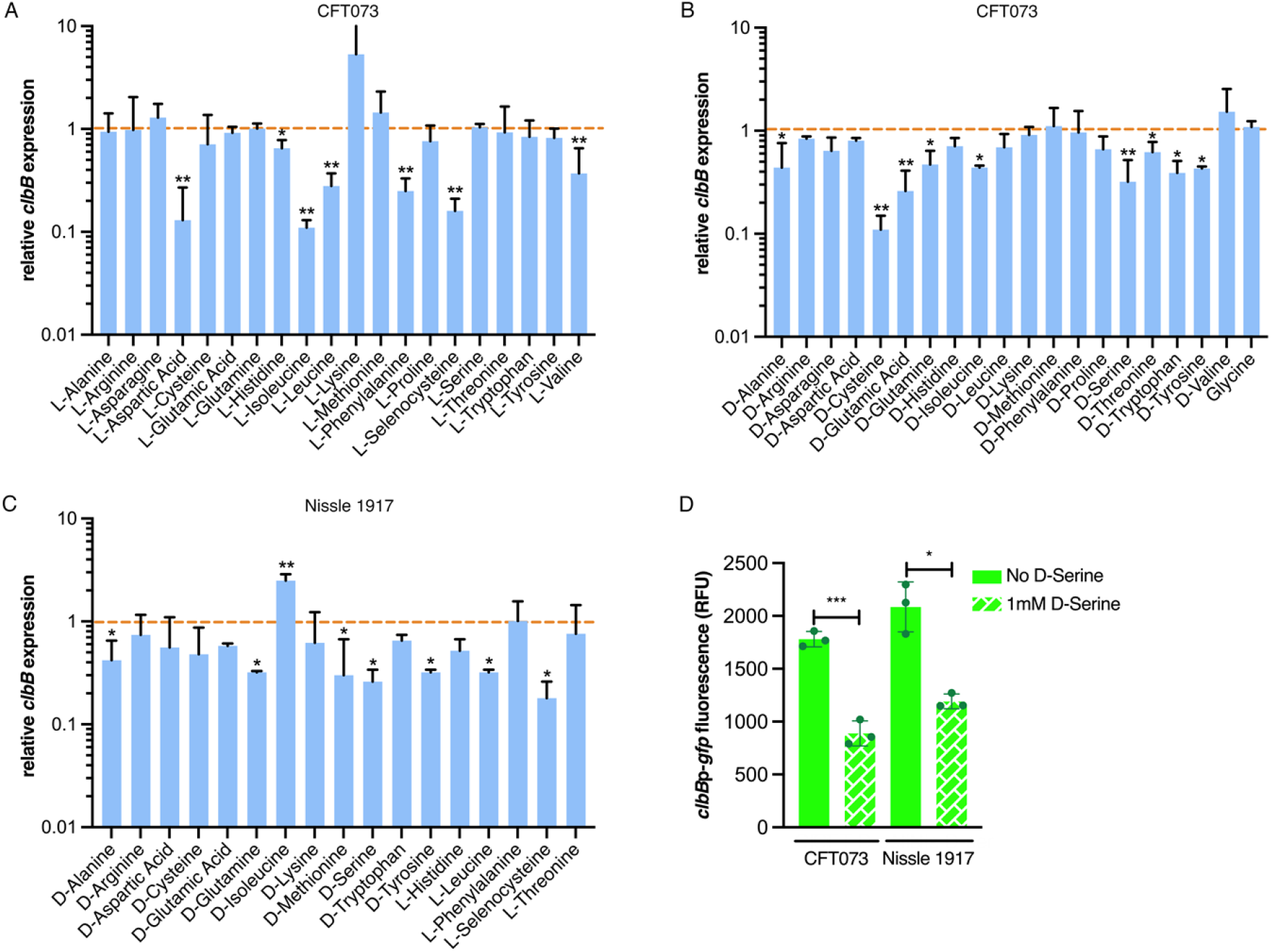
Expression of colibactin is modulated by L- and D-amino acids. Relative *clbB* expression was measured by RT-qPCR. CFT073 was grown in M9 minimal media supplemented with amino acids to a final concentration of 1 mM for 5 h. (**A**) Shows relative expression in the presence of L-amino acids and (**B**) indicates expression in the presence of D-amino acids. The orange dashed line indicates baseline expression with bars above and below this line representing up and downregulation, respectively. Statistical significance was determined from three biological replicates using an unpaired Student’s *t*-test with, * and ** indicating significance, *P* < 0.05 and 0.01, respectively. (**C**) Amino acids significantly affecting *clbB* expression in CFT073 were tested in Nissle 1917 under the same growth conditions as discussed above. (**D**) *clbBp*:gfp reporter activity in MEM-HEPES in the presence and absence of 1 mM D-serine. Bacteria were sampled at 4 h post-inoculation, *clbB* expression was measured as GFP/OD_600_. Columns represent mean +/− standard error of the means (SEM) with individual experimental observations indicated by data points. Statistical significance was assessed using a paired Student’s *t*-test with *, ***, denoting *P* < 0.05 and 0.0001, respectively.

### D-Serine induced repression of colibactin reduces DNA damage

Colibactin exerts a genotoxic effect in infected eukaryotic cells through the formation of DNA interstrand cross-links (ICLs) and DSBs [29]. As exposure to D-Serine caused a downregulation in the colibactin synthesis genes, we hypothesized that D-Serine would be capable of limiting the formation of DNA ICLs. To investigate this, we exposed linear DNA to live *pks*^+^ *E. coli* cultured in M9 minimal media alone or supplemented with 1 mM D-Serine and assessed colibactin-associated cross-linking activity. Upon exposure of linear DNA to Nissle 1917 and CFT073, strong cross-linking activity was apparent, as observed by an increase in molecular weight consistent with DNA duplex formation. However, upon the addition of D-Serine, a marked reduction in cross-linking activity was observed (Fig 3A). Quantification of DNA ICLs by densitometry revealed that the addition of D-Serine reduced the cross-linking of DNA by 2.83 and 1.39-fold, by Nissle 1917 and CFT073, respectively (Fig 3B). The formation of colibactin-associated ICLs in eukaryotic cells activates the ICL repair response pathway and results in the production of ICL-dependent DNA DSBs. Therefore, as exposure to D-Serine reduced cross-linking activity in linear DNA, we hypothesized that treatment with D-Serine could also lead to a reduction of DNA DSBs in HeLa cells infected with *pks*^+^ *E. coli*. To examine DSBs, we infected HeLa cells with Nissle 1917 and measured levels of γ-H2AX, a variant of the histone family H2A, that becomes phosphorylated in response to DNA DSBs. Cell lysates were extracted 4 h post-infection and detection of γ-H2AX was determined by immunoblotting (Fig 3C). To quantify the level of H2AX phosphorylation, normalized γ-H2AX signal intensities from three experiments were compared (Fig 3D). Nissle 1917 infected cells exhibited a 6.41-fold (*P* = 0.006) increase in normalized γ-H2AX signal compared with uninfected cells. Inclusion of D-Serine during infection with Nissle 1917 resulted in a 2.75-fold reduction in γ-H2AX signal intensity compared to cells infected with untreated Nissle 1917 (*P* = 0.003). Interestingly, D-Serine did not reduce γ-H2AX signal intensity upon infection with DH10B pBAC-*pks*, suggesting that D-Serine activity requires factors specific to the natural colibactin-carrying isolates described here. To confirm that D-Serine acted specifically on colibactin, DC10B was employed as a *pks^−^ E. coli* strain. γ-H2AX signal intensity indicated that levels of phosphorylation were similar to that of uninfected cells and the addition of D-Serine did not significantly change levels of phosphorylation. Taken together, these results indicate that D-Serine induced repression of natively encoded colibactin during infection of HeLa cells and led to a reduction in the formation of DSBs as measured by decreased phosphorylation of H2AX.

**Fig. 3.**
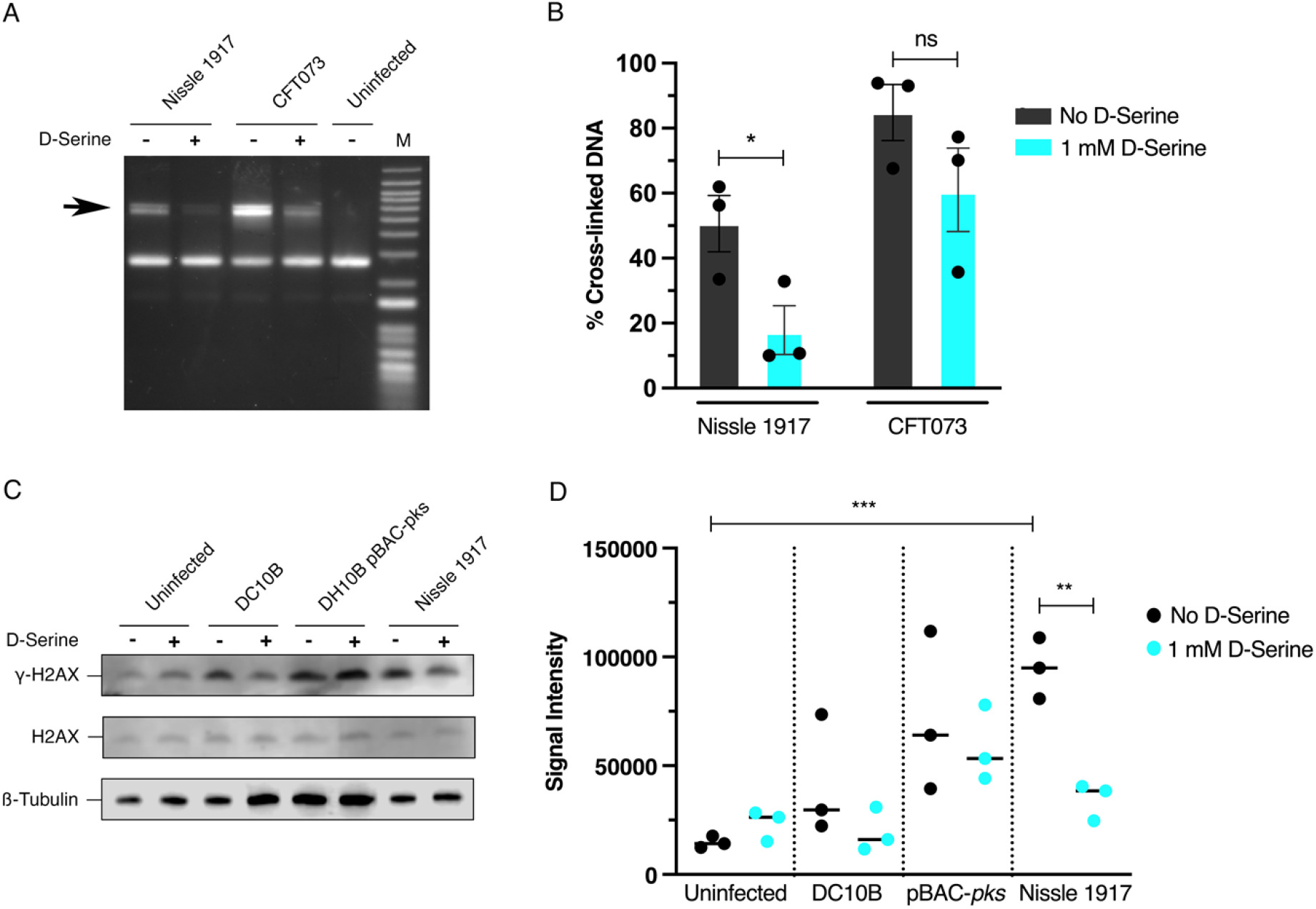
Repression of colibactin by D-Serine reduces cross-link formation, and DNA damage in Hela cell infection. (**A** and **B**) Nissle 1917 and CFT073 were cultured for 5 h in M9 minimal media alone (-) or in media supplemented with 1 mM D-Serine (+) before 1.5 × 10^6^ CFU was exposed to linearized plasmid DNA for 40 min. The DNA was extracted, and cross-linking activity was determined by electrophoresis in denaturing conditions. (**A**) DNA cross-link formation of linearized plasmid DNA exposed to Nissle 1917 and CFT073 was visualized after migration under alkaline denaturing conditions. M, DNA size marker (1 kB plus DNA ladder, Invitrogen). (**B**) The percentage of the DNA signal in the upper, cross-linked band (indicated by the arrow in panel **A**) relative to the total DNA signal. Signal intensities were quantified using ImageJ for three independent experiments and statistical significance was assessed by unpaired Student’s *t*-test with, * indicating *P* = < 0.01. (**C** and **D**) HeLa cells were infected for 4 h with live *pks*^+^ and *pks*^−^ *E. coli* with a multiplicity of infection (MOI) of 400 bacteria per cell or left uninfected. Infections were performed in wells containing MEM-HEPES alone (-) or with media supplemented with 1 mM D-Serine (+). (**C**) Immunoblot analysis of cell lysates extracted 4 h post infection. Phosphorylated histone (γ-H2AX) was used as an indicator of double stranded DNA breaks and total histone (H2AX) was used as an internal control. Β-Tubulin was used as a loading control for cell lysates. DH10B pBAC*-pks* and DC10B were used as positive and negative controls, respectively. (**D**) Signal intensities of bands were measured using LI-COR Image Studio software. γ-H2AX signals were corrected to account for any variation in loading using β-Tubulin signal intensity. Experimental signal was normalized so that the mean signal intensity of the eight samples was equivalent for each experiment. The experiment was carried out in triplicate. Columns represent mean +/− SEM with individual experimental observations indicated by data points. Statistical significance was assessed by unpaired Student’s *t*-test with, ** and *** indicating *P* < 0.01 and 0.001, respectively.

### Double strand break lesion formation is reduced upon exposure *of pks*+ *E. coli* to D-Serine

In response to DNA DSBs, factors involved in the DNA damage response (DDR), including γ-H2AX, accumulate at sites of damage temporarily [52]. These so-called nuclear foci are formed by γ-H2AX which can spread over megabases along the DNA flanking the breakage site [53] and can be visualized using specific antibodies under a fluorescent microscope [54]. Therefore, we investigated the effects of D-Serine on the formation of DNA DSBs at the subcellular level by visualizing nuclear foci positive for γ-H2AX [55]. HeLa cells were infected with *pks*^+^ *E. coli* and then labelled with anti-γ-H2AX monoclonal antibody before visualizing by confocal microscopy. We observed intense punctate staining with anti-γ-H2AX after infecting cells with untreated Nissle 1917 indicative of the induction of DNA DSBs. However, exposure to D-Serine led to the formation of fewer nuclear lesions/foci. The nuclei resembled those of uninfected cells, and displayed few γ-H2AX foci, whereas the number of foci remained unchanged in cells infected with DH10B pBAC-*pks* both in the presence and absence of D-Serine (Fig 4A). To investigate the heterogeneity in the response of individual HeLa cells to D-Serine during infection γ-H2AX phosphorylation was measured by flow cytometry. This revealed that D-Serine reduced the number of γ-H2AX positive cells by 2.84-fold. However, exposure to D-Serine did not fully return levels of γ-H2AX fluorescence to that of the uninfected cells, (Fig 4B).

**Fig 4.**
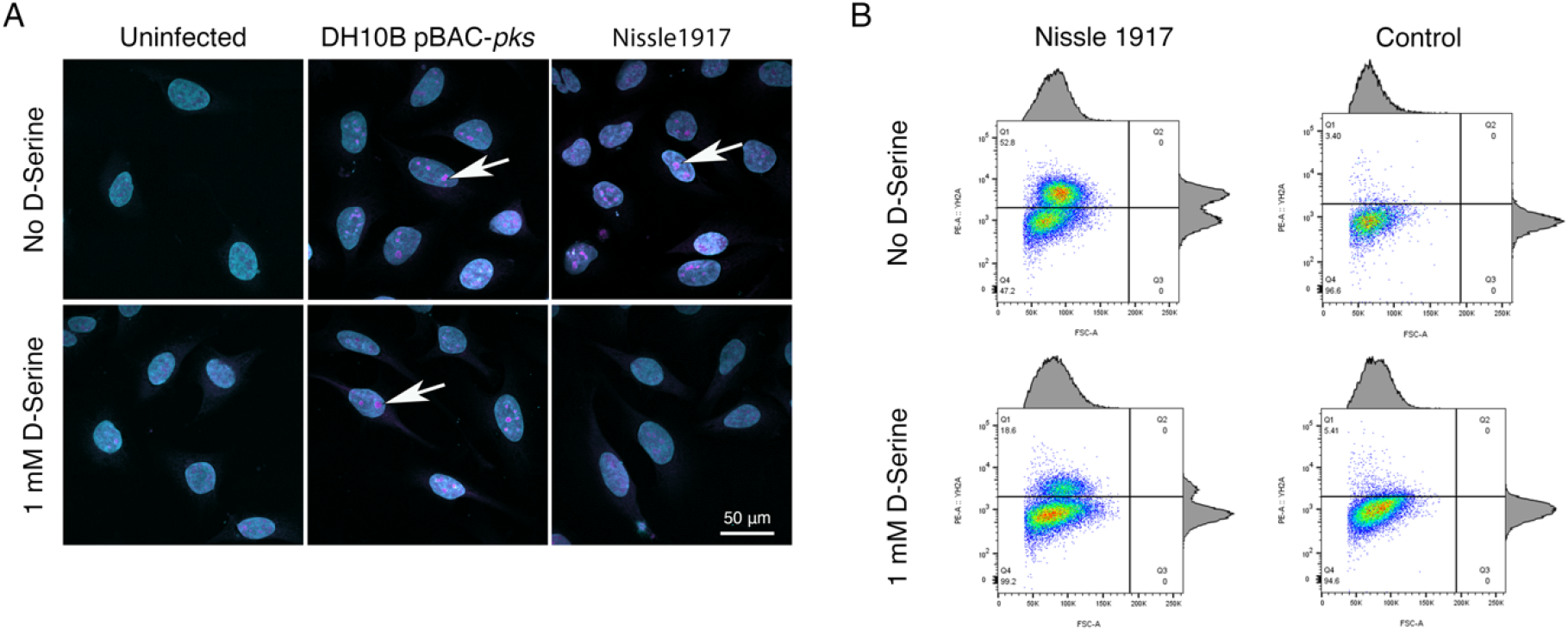
D-Serine reduces nuclear foci observed in HeLa cells. HeLa cells were infected for 4 h with *E. coli* Nissle 1917 or DH10B hosting BAC-*pks* (MOI = 400). Infections were performed with and without the addition of 1 mM D-Serine to the growth media. At 8 h post infection, cells were washed, fixed, and stained with anti-γ-H2AX antibody. (**A**) Cells were examined by confocal microscopy for DNA in cyan and phosphorylated histone H2AX protein in magenta. Images of uninfected, and *pks*^+^ infected cells are shown, scale bar = 50 μm. (**B**) Intracellular levels of phosphorylated histone H2AX were measured by flow cytometry 8 h after infection. Dot plots reveal the percentage of viable cells fluorescing in the γ-H2AX channel, 100k events were analysed for each sample.

### Exposure to D-Serine reduces colibactin-associated cellular senescence

Cellular senescence has been described as an irreversible state of cell-cycle arrest, often in response to DNA damage [56]. This phenomenon is associated with colibactin and is observed when mammalian cells are infected with live *pks^+^ E. coli* [22]. Exposure to D-Serine led to a reduction in H2AX phosphorylation, demonstrating that colibactin-associated DNA damage was reduced by D-Serine (Fig 4). Therefore, we aimed to show that exposure to D-Serine would also result in a reduction in downstream cellular senescence. HeLa cells were infected with live *pks^+^ E. coli* for 4 h, before cells were treated with gentamicin for 72 h. Cell morphology was observed by staining actin filaments with Phalloidin-Alexa Fluor 555 and using fluorescence microscopy for visualization (Fig 5A). Colibactin-producing *E. coli* have been distinguished by their ability to induce megalocytosis in cultured eukaryotic cells, a phenotype associated with senescence, characterized by progressive enlargement of the cell body, and the nucleus, and the abolishment of mitosis (23). In the absence of D-Serine, cells infected with Nissle 1917 displayed the characteristic cytopathic effect, with cellular and nuclear enlargement being apparent (Fig. 5). However, upon exposure to D-Serine, cells infected with Nissle 1917 had cell morphology that was markedly more like that of the uninfected control. Consistent with our findings described earlier (Fig 3), D-Serine treatment did not result in a decrease in the senescence-associated morphological alterations observed in DH10B pBAC-*pks*. The mean cell area was determined for three independent replicates using image analysis software, CellProfiler (Fig 5B). Cell enlargement increased 2.60-fold (from 1209.52 to 3140.90 μm) during infection with Nissle 1917, however, upon addition of D-Serine, the mean cell area decreased to 666.62 μm (*P* < 0.01). Individual cell area measurements indicate the variable extent of cell enlargement observed across the three experiments (Fig 5C). Overall, these results demonstrated that reduction of colibactin expression through exposure to D-Serine reduced both the acute and more long-term colibactin-related cytopathic effects of infection with Nissle 1917.

**Fig 5.**
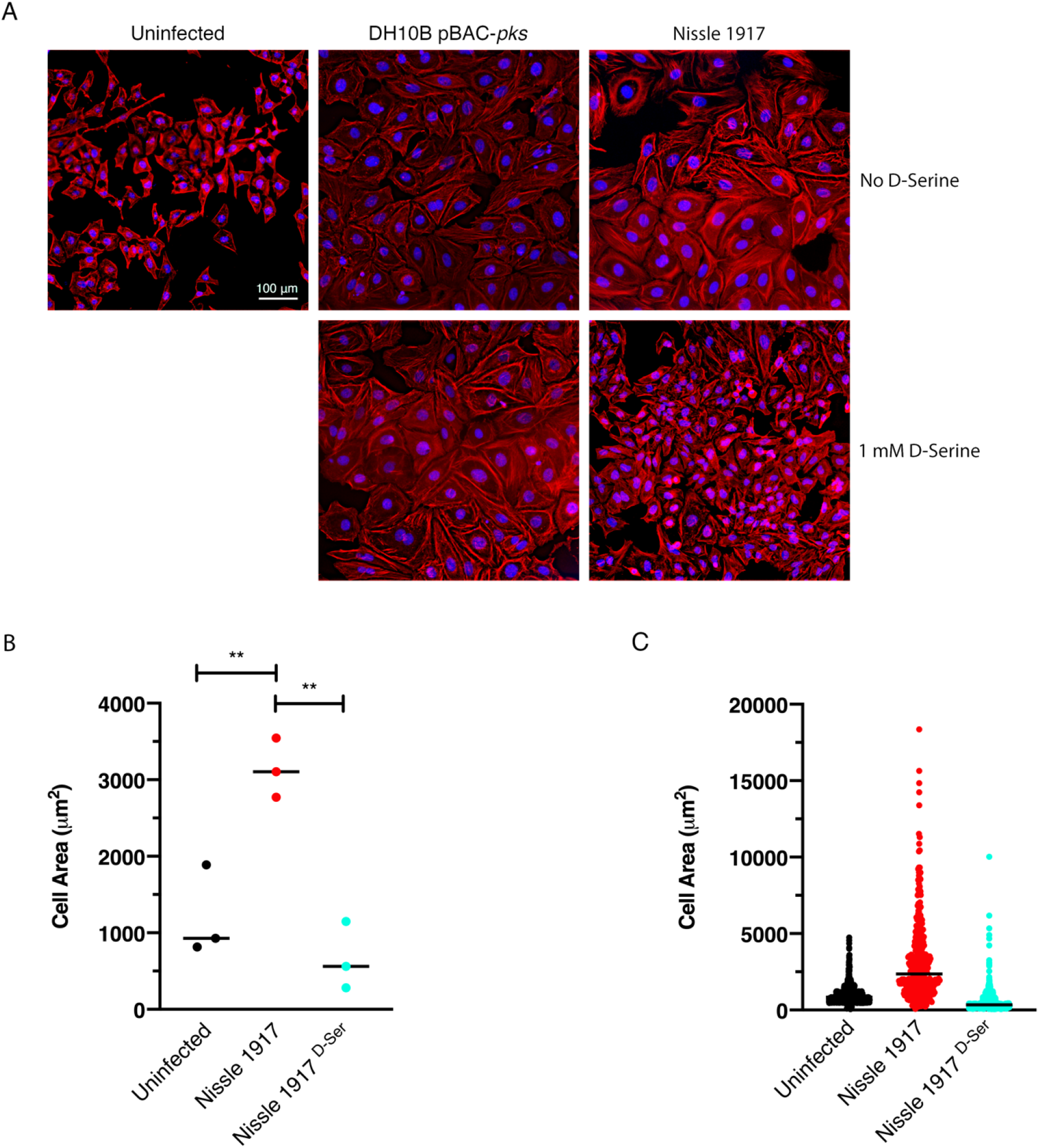
Exposure to D-Serine reduces the colibactin associated cellular senescence. HeLa cells were infected for 4 h with *E. coli* Nissle 1917 or DH10B hosting BAC-*pks* (MOI = 400). Infections were performed with and without the addition of 1 mM D-Serine to the growth media. At 8 h after infection, cells were washed and incubated for 72 h to allow for the megacell phenotype to develop. (**A**) HeLa cell morphology was observed by wide field fluorescence. Actin cytoskeleton was stained with Phalloidin in red and DNA was counterstained with DAPI in blue at 72 h post infection. Scale bar = 100 μm. CellProfiler software was employed to measure the area of HeLa cells using images acquired at 10X magnification. 100 cells were measured per image. (**B**) Columns represent the mean cell area measured with individual experimental observations indicated by data points for each infection condition. Measurements were acquired from images taken from three independent experiments and statistical significance was assessed by unpaired Student’s *t*-test with, ** indicating *P* < 0.01. (**C**) Individual cell area measurements were recorded across triplicate experiments. Black lines indicate the mean.

### The D-Serine metabolism locus is not essential for repression of colibactin in *pks*+ *E. coli* in response to D-Serine

DsdC is a well-studied D-Serine responsive transcriptional regulator that is required for catabolism of D-Serine via activation of *dsdXA* [16]. We hypothesized that the repression of the *pks* locus by D-Serine might be mediated by DsdC. Therefore, *dsdC* was deleted in Nissle 1917 and the effect on colibactin production with and without D-Serine was assessed. First, we compared transcriptomes from a previous study [21], which revealed that exposure to D-Serine triggered similar reductions in *clb* gene expression in both WT and ∆*dsdC* CFT073 strains (Fig 6A). To assess the effects on genotoxic activity, HeLa cells were infected with Nissle ∆*dsdC* as described previously. Inclusion of D-Serine resulted in a 6.34-fold reduction in *clbB* expression compared to cells infected with Nissle ∆*dsdC* alone (*P* = 0.0145; Fig. 6B and C). Furthermore, HeLa cells infected with Nissle ∆*dsdC* in the presence of D-Serine were protected from genotoxic attack as lower levels of phosphorylated H2AX were detected, indicating that the DNA repair response was reduced in both wild type and ∆*dsdC* genetic backgrounds. Taken together, these data demonstrate that D-Serine-associated downregulation of *pks* encoded genes occurs independently of DsdC.

**Fig 6.**
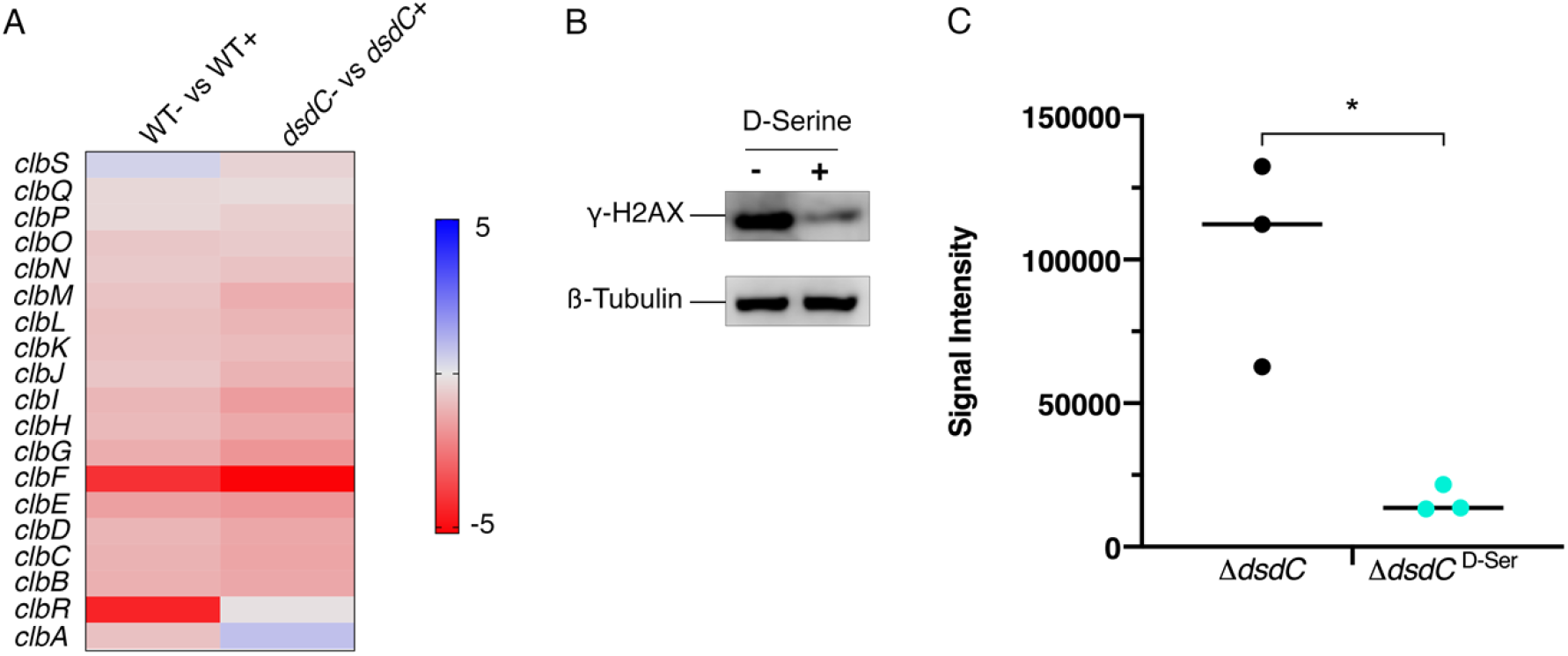
Deletion of *dsdC* does not affect D-Serine associated repression of colibactin. Generation of an isogenic mutant, Nissle ∆*dsdC,* revealed that D-Serine-associated repression of colibactin activity occurred independently of the D-Serine tolerance locus in *pks*^+^ *E. coli* strains. (**A**) Heat map showing the EdgeR calculated log_2_ relative fold changes for each gene in the *pks* island with corresponding colour key adjacent. False discovery rate-corrected *P* values can be found in Table S3 (**B**) The genotoxic activity of Nissle ∆*dsdC* was assessed by infecting HeLa cells as discussed above. Proteins were extracted and the level of H2AX phosphorylation was determined. Immunoblot analysis of cell lysates extracted 4 h post-infection is shown. Anti-γ-H2AX antibody was used as an indicator of double stranded DNA breaks and β-Tubulin was used as a loading control for cell lysates. (**C**) Signal intensities of bands were measured as described in the methods using LI-COR Image Studio. γ-H2AX signals were corrected to account for any variation in loading using β-Tubulin signal intensity. Experimental signal was normalized so that the mean signal intensity of the samples was equivalent for each experiment. The experiment was carried out in triplicate. Columns represent mean +/− SEM with individual experimental observations indicated by data points. Statistical significance was assessed by unpaired Student’s *t*-test with, * indicating *P* < 0.05.

## Discussion

Sensing and responding to environmental cues and signalling molecules is crucial for bacterial survival. Indeed, modulating gene expression enables bacteria to adapt and persist in changing environments; a trait that is essential for certain *E. coli* pathotypes that can colonize multiple sites of the human host. For instance, UPEC strains can survive in both the gastrointestinal tract and the bladder. Strikingly, previous work has shown that UPEC strains extensively carry the *dsdCXA* locus [10], enabling these strains to metabolize D-Serine in the nutrient deficient bladder. Furthermore, metabolizing D-Serine has been shown to confer a fitness advantage in strains that infect this site [9]. Indeed, there has been a growing appreciation for the important role of amino acids as carbon sources; however, there has also been an emphasis on investigating the regulatory functions of certain amino acids. Investigations into *E. coli* biofilm formation revealed the spatiotemporal regulation of L-Alanine metabolism is essential for cell viability and growth of colonies [57], whereas conversely, catalysis of the amino acid L-Tryptophan was implicated with the inhibition of biofilm formation [58]. In the mammalian host, sensing these metabolites can serve as stimuli to trigger the expression of essential virulence genes. In response to L-Arginine in the gut, pathogenic EHEC strains signal the upregulation of LEE-encoded genes to facilitate site-specific colonization of the host [59]. However, in contrast, exposure to D-Serine has been implicated in the downregulation of virulence genes in this *E. coli* pathotype, with our previous work demonstrating that D-Serine represses the type three secretion system (T3SS) [10]. While our work has shown that D-Serine is present in trace concentrations in the gut [10], approximately 1000-fold lower than the concentration reported for the bladder [9], the production of D-Serine by members of the gut microbiome has been reported [60]. As a result, the *E. coli* strains residing in the gut may encounter localized micro-niches rich in this metabolite, raising the possibility that it functions as a niche-specific regulator of diverse virulence genes in pathogenic *E. coli*. In this study we observed that, in general, D-amino acids exerted an enhanced ability to regulate expression of the genes involved in colibactin production over their L-amino acid enantiomers. We demonstrated that treatment with several D-amino acids, most notably D-Cysteine and D-Serine, resulted in downregulation of the *pks* genomic island in colibactin producing *E. coli.* Interestingly, analysis of gene expression in CFT073, revealed that L-Cysteine and L-Serine, did not significantly repress *clbB*. Therefore, while L-amino acids are favoured in nature [3], D-amino acids can have distinct effects in regulating gene expression and the mechanisms underpinning these effects warrant further investigation.

Colibactin research has been invigorated after a recent breakthrough study revealed that exposure to colibactin caused a specific mutational signature that linked *pks*^+^ harbouring *E. coli* to CRC tumours [36]. Thus, researchers have endeavoured to identify compounds that will inhibit colibactin and prevent the pro-tumorigenic effects. Cougnoux *et al*., (2016) described the use of boron-based compounds that functioned as enzyme competitors and inhibited the activity of ClbP, the serine peptidase involved in colibactin maturation [61]. Furthermore, the use of these compounds prevented the genotoxic and tumorigenic activity of colibactin on epithelial cells and in a CRC mouse model [61]. Similarly, mesalamine – an anti-inflammatory drug that is used for treating inflammatory bowel disease (IBD) and is associated with reduced risk of CRC in IBD patients [62] – has been shown to reduce *clbB* expression and hence inhibit production of colibactin [63]. The increasing prevalence of drug resistance has led to the development of new and natural antimicrobial agents, some of which have interesting biological activities beyond their intended use as bactericidal agents. For example, cinnamon and its essential oil (cinnamaldehyde) have been studied for their antibacterial properties [64]. Interestingly, recent research showed that treatment with these compounds induced downregulation of the *pks*-encoded *clbB* gene *in E. coli* strains isolated from patients with CRC [65]. In addition, tannin, a compound extracted from medicinal plants, was also shown to repress transcription of colibactin and prevent the associated genotoxic activity of colibactin producing *E. coli* [66]. In this study we have highlighted the important role of naturally available amino acids, by identifying D-Serine, as a potent repressor of colibactin. We showed that exposure to D-Serine prevented the colibactin-associated cytopathic effects in eukaryotic cells and that treatment with 1 mM D-Serine was sufficient to induce prolonged protection, as signs of cellular senescence remained absent after 72 h. This suggests that D-Serine could have prophylactic potential, providing the host with long-lived protection against colibactin production by commensal *E. coli* residing in the gastrointestinal tract. Such a treatment would particularly benefit high-risk patients, such as individuals with IBD, where the prevalence of *E. coli* belonging to the B2 phylogroup is high and the incidence of developing CRC is significantly greater [67,68]. While our results indicate promising potential for D-Serine as a therapeutic, it should be noted that colibactin has been detected in urine (an environment rich in D-Serine [9]) from individuals with *pks*+ urinary tract infections [69]. In the same study, colibactin-induced DNA damage was observed in a murine model of cystitis. Levels of D-Serine can vary across individuals and be affected by diet, hence further work will be needed to understand the potential of D-Serine to repress colibactin in complex host environments.

Bacterial gene regulatory networks tightly govern the expression of genes in response to chemical or environmental stimuli; however, current understanding on the regulation of the colibactin gene locus is limited. Homburg *et al.,* (2007) sought to elucidate the transcriptional organization and regulation of the colibactin genes using Nissle 1917 as a model [44]. Their results showed that the *clb* locus could be divided in to at least seven transcriptional units, of which four were found to be transcribed polycistronically. The polycistronic expression of *clbR*/*clbA* indicated a potential regulatory function exerted by the *clbR* encoded LuxR-like regulatory protein on *clbA.* Indeed, the expression of *clbA* is crucial for colibactin production, as it encodes a phosphopantetheinyl transferase that is responsible for the post-translational activation of the PKS and NRPS proteins of the colibactin biosynthesis pathway [44]. Recently, *clbR* was identified to encode the key transcriptional activator of the *clb* genes, and expression of this gene was found to directly correlate with the function and production of colibactin in *E. coli pks*^+^ strain M1/5 [70]. Different carbon sources have been found to influence *clbRA* transcript levels, with *clbA* and *clbR* upregulated in the presence of glucose and glycerol compared with pyruvate and acetate, where expression levels remained at an intermediate level [44]. Increased expression of *clbA* was found to be associated with exposure to nondigestible oligosaccharides commonly found in prebiotics such as lactose and raffinose [71], inulin and galacto-oligosaccharides [72]. Inulin also increased expression of *clbB*, *clbQ* and *clbR* [72]. These data highlight that the diet may play a significant role in controlling the production of colibactin, with certain foods potentially creating a pro-tumorigenic environment. Interestingly, iron homeostasis has also been implicated with the production of colibactin, with *clbA* downregulated and colibactin-associated megalocytosis reduced upon exposure to FeCl_3_ [73]. Furthermore, supplementation of iron into media containing colibactin-inducing oligosaccharides resulted in the abrogation of *clbA* induction [72]. The production of colibactin was shown to be regulated by the ferric uptake regulator (Fur) by Tronnet *et al.*, (2016). The study revealed that Fur positively regulated *clbA* expression, and that transcription was initiated by direct binding of Fur to the *clbA* promoter [74]. However, iron-dependent transcription of *clbA* was found to be independent of *clbR* [74], suggesting the involvement of a second *clbA* promoter that may be activated upon specific iron conditions. Iron availability is tightly controlled in the host, and Gram-negative bacteria have evolved several iron uptake systems to overcome this limitation, including the secretion of iron sequestering siderophores [75]. Intriguingly, *clbA* has been shown to have diverse functionality, contributing to the synthesis of both colibactin and the yersiniabactin siderophore [76]. Taken together, these reports suggest that complex regulatory networks are involved in the production of colibactin and that transcription of *clb* genes may be activated under specific nutritional conditions which could serve as a fitness advantage to *pks*-harbouring bacteria.

D-Serine exposure has been demonstrated to induce the activation of the LTTR DsdC, whose main function has been described to be the activation of *dsdXA* transcription [16] in response to D-Serine. This cluster facilitates uptake and catabolism of D-Serine, respectively. As DsdC is the primary transcriptional regulator induced in response to D-Serine, we investigated whether the repression of *clb* genes was mediated by DsdC. Our data indicate that repression of colibactin by D-Serine is not directly facilitated by DsdC, as similar responses to D-Serine were observed in both WT and Nissle Δ*dsdC* both in terms of repression of *clb* gene expression and cytopathic responses. These findings were unexpected. Interestingly, Wallenstein *et al.*, (2020) recently demonstrated that the *pks*-encoded *clbR* gene is the transcriptional activator of the colibactin genes. Like DsdC, ClbR has been described as a LTTR with a helix-turn-helix DNA-binding motif that interacts with the *clbR*-*clbB* intergenic region, suggesting it is involved in the regulation of the *pks* island [44,70]. LTTR regulators are thought to be activated by a small molecule co-inducer [77]. Therefore, it is also plausible to consider that D-Serine could enter the cell by an alternative transport system and act as a co-inducer of ClbR, Fur, or indeed another elusive LTTR. Indeed, D-Serine can enter the cell via CycA [78,79] thereby potentially explaining why exposure to D-Serine reduced *clbB* expression in both Nissle Δ*dsdC* and in the WT. The precise regulatory mechanism by which D-Serine causes a reduction in colibactin gene expression remains unsolved and will be a focus of future work in our group. In conclusion, in this study we have identified a D-amino acid that has a strong regulatory effect on the genes encoded in the *pks* genomic island. Exposure to D-Serine in *pks*^+^ *E. coli* results in downregulation of the *clb* genes and consequently, reduced DNA cross-linking and a reduction in the phenotypic responses associated with colibactin-induced DNA damage in cultured eukaryotic cells. Furthermore, deletion of *dsdC* revealed that D-Serine-induced inhibition of colibactin is not mediated by the D-Serine metabolism locus regulator, DsdC. Understanding the physiological implications *in vivo* will be key in further exploring the prophylactic potential of D-Serine.

## Acknowledgements

We thank Diane Vaughan and Alana Hamilton (Flow Core Facility, University of Glasgow, United Kingdom) for their assistance with processing samples for flow cytometry and Dr Leandro Lemgruber Soares and Susan Baillie (Glasgow Imaging Facility, University of Glasgow, United Kingdom) for their assistance with the acquisition of confocal images. We are very grateful to Dr Michael Ormsby (University of Stirling, United Kingdom) and Dr James Connolly (Newcastle University, United Kingdom) for their insightful appraisal of our manuscript. This work is supported by the Biotechnology and Biological Sciences Research Council, grant numbers BB/M029646/1 and BB/R006539/1. The work performed in Toulouse, France, was funded by the French National Agency for Research (ANR), grant number UTI-TOUL ANR-17-CE35-0010 and ANR-19-AMRB-0008.

## Supporting information

**S1 Table. Strains and plasmids used in this study.**

**S2 Table. Oligonucleotide sequences used in this study.**

**S3 Table. Fold changes and *P* values of *clb* genes in CFT073 determined by RNA-Seq**

**S4 Table. Relative gene expression and *P* values in CFT073 determined by RT-qPCR**

**S5 Table. Relative gene expression and *P* values in Nissle 1917 determined by RT-qPCR**

